# Spatial Organization of Neuron-Astrocyte Interactions in the Somatosensory Cortex

**DOI:** 10.1101/2022.02.03.479046

**Authors:** Andrés M. Baraibar, Lindsey Belisle, Giovanni Marsicano, Carlos Matute, Susana Mato, Alfonso Araque, Paulo Kofuji

**Affiliations:** Department of Neuroscience, University of Minnesota. Minneapolis, USA; INSERM, U862 NeuroCentre Magendie, Endocannabinoids and Neuroadaptation, Bordeaux, France; University of Bordeaux, NeuroCentre Magendie, Endocannabinoids and Neuroadaptation, Bordeaux, France; Department of Neurosciences, University of the Basque Country UPV/EHU, Leioa, Spain; Achucarro Basque Center for Neuroscience, Leioa, Spain; Biocruces Bizkaia, Baracaldo, Spain; Centro de Investigación Biomédica en Red sobre Enfermedades Neurodegenerativas (CIBERNED), Madrid, Spain

## Abstract

Microcircuits in the neocortex are functionally organized along layers and columns, which are the fundamental modules of cortical information processing. While the function of cortical microcircuits has focused on neuronal elements, much less is known about the functional organization of astrocytes and their bidirectional interaction with neurons. Here we show that CB_1_R-mediated astrocyte activation by neuron-released endocannabinoids elevate astrocyte Ca^2+^ levels, stimulate ATP/adenosine release as gliotransmitters, and transiently depress synaptic transmission in layer 5 pyramidal neurons at relatively distant synapses (>20 µm) from the stimulated neuron. This astrocyte-mediated heteroneuronal synaptic depression occurred between pyramidal neurons within a cortical column and was absent in neurons belonging to adjacent cortical columns. Moreover, this form of heteroneuronal synaptic depression occurs between neurons located in particular layers, following a specific connectivity pattern that depends on a layer-specific neuron-to-astrocyte signaling. These results unravel the existence of astrocyte-mediated non-synaptic communication between cortical neurons, and that this communication is column- and layer-specific, which adds further complexity to the intercellular signaling processes in the cortex.

## INTRODUCTION

The neocortex is the most complex structure of the mammalian brain involved in higher cognitive functions. The cellular organization of the cerebral cortex is well known since the work of Cajal and his disciple Lorente de Nó, who proposed that cortical neurons form functional modules that serve as the “elementary cortical unit of operation”^1,2^. Cortical neurons are organized horizontally in six layers and vertically in columns^3^. The columnar configuration of the neocortex is a widely accepted idea explaining its functional organization^4,5^. A great amount of information has been provided regarding the neuronal elements involved in cortical circuits and their synaptic microorganization^6^. However, the properties of non-neuronal cell types, like astrocytes, and their functional interactions with neurons in this elementary module remain largely unexplored.

Astrocytes have emerged as key regulatory elements of synapses, responding with Ca^2+^ elevations to synaptically-released neurotransmitters and releasing gliotransmitters that regulate synaptic transmission in different brain areas^7–9^. In the cortex, sensory stimuli or direct neuronal stimulation evoke astrocyte Ca^2+^ elevations^10–17^, which are topographically represented in the primary somatosensory cortex S1^18^ and spatially restricted to the cortical columns in the barrel cortex^14^. Cortical astrocyte Ca^2+^ elevations can, in turn, stimulate the release of gliotransmitters, such as glutamate or D-Serine, that can regulate synaptic transmission^13,15^, and that can be responsible for the observed astrocyte-mediated regulation of the cortical network function^17,19–21^. Moreover, synaptic regulation by astrocytes may be exerted at synapses relatively distant from the active synapses^22–24^. This phenomenon, termed lateral astrocyte synaptic regulation^25^, resembles the classical heterosynaptic modulation but is mechanistically dissimilar because it involves astrocytes and may be crucial in brain circuits where spatial signaling greatly influences neural network function, like the neocortical columns. However, the spatial properties of astrocyte-neuron interaction and the consequent synaptic regulation in defined cortical columns and layers remain unidentified.

Endocannabinoid (eCB) signaling has been proposed to mediate astrocyte-neuron communication in different brain regions, including the neocortex. In particular, endogenous activation of astroglial type-1 cannabinoid receptors (CB1Rs) regulates hippocampal and neocortical synaptic transmission and plasticity^15,22,26,27^. Here, in order to decipher the regulatory role of astrocytes in the synaptic physiology of cortical columns, we took advantage of this eCB signaling to physiologically stimulate cortical astrocytes. We show that eCBs released from layer 5 (L5) pyramidal neurons induce Ca^2+^ elevations in astrocytes and transiently depressed synaptic transmission in adjacent pyramidal neurons. This form of heteroneuronal synaptic depression required astrocytic cannabinoid receptor type (CB_1_R) activation and was mediated by presynaptic type 1 adenosine receptors (A1Rs). Astrocyte-mediated heteroneuronal synaptic depression was present between pyramidal neurons within a cortical column and was absent in neurons belonging to adjacent cortical columns. Moreover, this form of heteroneuronal synaptic depression occurred between neurons located in particular layers, following a specific connectivity pattern that depends on a layer-specific neuron-to-astrocyte signaling. These results reveal the existence of astrocyte-mediated non-synaptic communication between cortical neurons, which is column- and layer-specific, and which adds further complexity to the intercellular signaling processes in the cortex.

## RESULTS

### Endocannabinoid signaling induces homoneuronal and heteroneuronal synaptic depression in S1

To investigate the spatial regulation of synaptic transmission in the primary somatosensory cortex, we performed double patch-recordings of layer 5 (L5) pyramidal neurons and monitored excitatory postsynaptic currents (EPSCs) evoked by electrical stimulation of layer 2/3 (L2/3). We then stimulated one neuron by a depolarizing pulse and recorded synaptic currents in both the “stimulated” neuron (homoneuronal synapses) and the adjacent (70-270 µm apart) “nonstimulated” neuron (heteroneuronal synapses) (**Figures 1A and 1B)**. Stimulation of single L5 pyramidal neurons induced a transient synaptic depression in 36 out of 93 (38.7%) homoneuronal synapses. Furthermore, in simultaneously recorded heteroneuronal synapses, this neuronal depolarization (ND) also induced a transient depression of synaptic transmission in 20 out of 72 (27.8%) heteroneuronal synapses (**Figures 1C-E**). Both homoneuronal and heteroneuronal synaptic depressions could be reliably induced by repeated stimulations (**Figure S1A and S1B**) and were associated with changes in the paired-pulse ratio (PPR), which are consistent with presynaptic mechanisms (**Figures S1C and S1D**).

**Figure 1.**
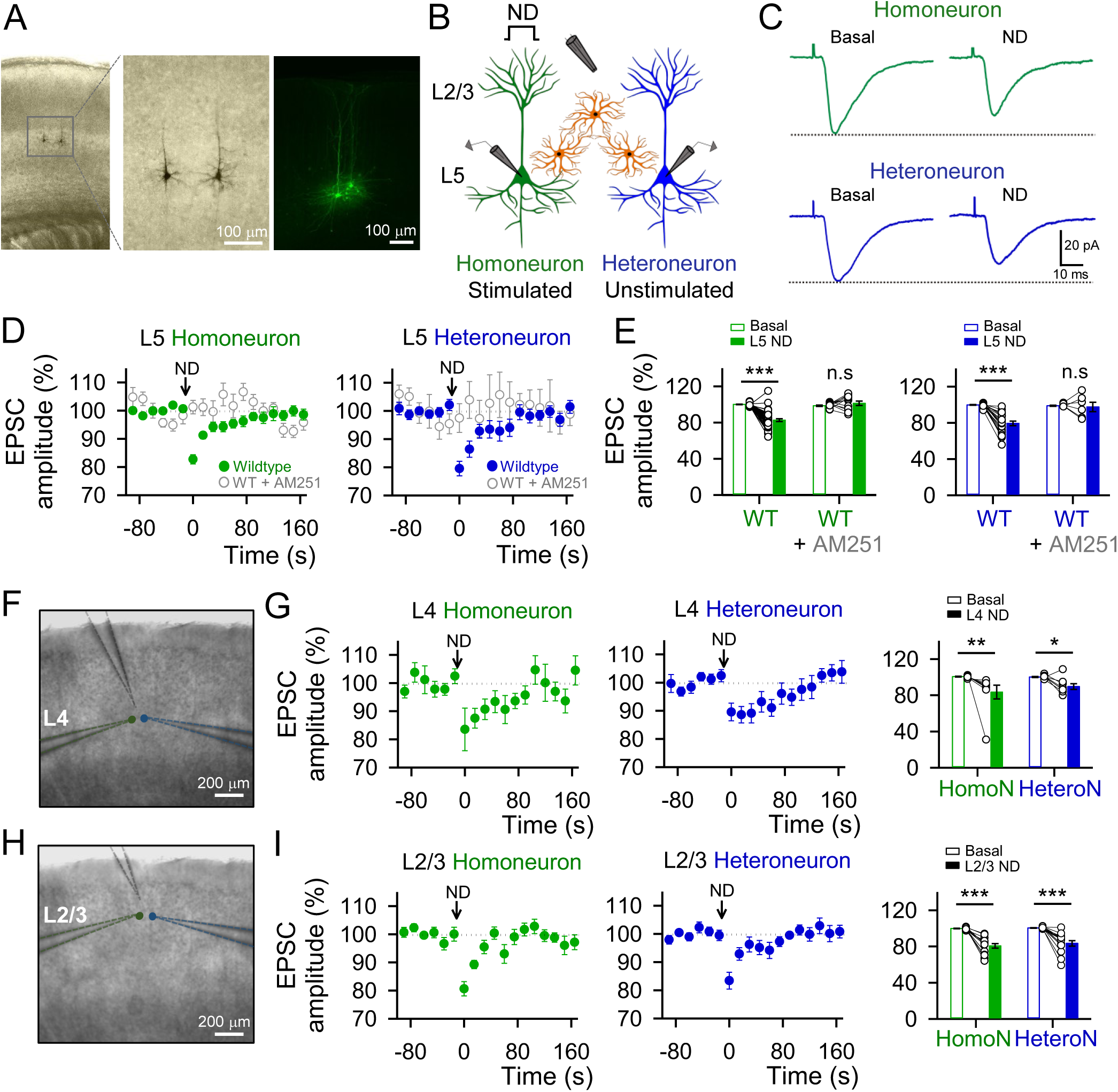
Endocannabinoid signaling induces homoneuronal and heteroneuronal synaptic depression in S1. (**A**) Biocytin loading S1 L5 pyramidal neurons image. (**B**) Schematic drawing depicting double patch-recordings from L5 pyramidal neurons and the stimulating electrode in L2/3. (**C**) Averaged EPSCs (n = 20 stimuli) before (control) and after neuronal depolarization (ND) in wildtype mice. (**D**) EPSCs amplitude *versus* time before (basal) and after ND in control (green or blue) and in the presence of AM251 (2 µM; open grey) in the homoneuron (left) and heteroneuron (right) from layer 5. (**E**) Relative changes in EPSC amplitude in control and with AM251 (2 µM). Two-tailed Student’s paired t test. (**F**) Representative infrared differential interference contrast image of the experimental configuration with the stimulation pipette in layer 4 and the homoneuronal and heteroneuronal neurons located in layer 4. (**G**) EPSCs amplitude *versus* time before (basal) and after ND in the homoneuron (left, green) and heteroneuron (middle, blue) in the experimental conditions represented in panel **F**. Right: Relative changes in EPSC amplitude. (**H**) Representative infrared differential interference contrast image of the experimental configuration with the stimulation pipette in L2/3 and the homoneuronal and heteroneuronal neurons located in L2/3. (**I**) EPSCs amplitude *versus* time before (basal) and after ND in the homoneuron (left, green) and heteroneuron (middle, blue) in the experimental conditions represented in panel **H**. Right: Relative changes in EPSC amplitude. Two-tailed Student’s paired t test. Data are expressed as mean ± SEM, *p < 0.05, **p < 0.01, ***p < 0.001.

ND is known to trigger the release of eCBs^28,29^ that can directly affect relatively close synapses (∼20 μm)^29–32^, a phenomenon called depolarization-induced suppression of excitation (DSE)^29–34^, and indirectly regulate more distant synapses through stimulation of astrocytes^15,22,27^, a phenomenon called astrocyte-mediated lateral regulation of synaptic transmission. Consistent with eCB-mediated synaptic regulation, homoneuronal and heteroneuronal synaptic depressions observed under control conditions were abolished following bath perfusion with the cannabinoid receptor type 1 (CB_1_R) antagonist AM251 (2 μM; n = 12 and 6), indicating that both phenomena were mediated by CB_1_R activation (**Figures 1D and 1E**).

We then tested whether these phenomena were present in other cortical layers by performing paired recordings of neurons in L2/3 and L4. L2/3 or L4 ND induced both homoneuronal (14 out of 38 and 8 of 32 pairs; 36.8% and 25%, respectively) and heteroneuronal (14 out of 39 and 8 of 27 pairs; 35.9% and 29.6%, respectively) depression (**Figures 1F-I**), indicating that eCB-induced homoneuronal and heteroneuronal synaptic depressions are a general cortical phenomena.

### Heteroneuronal, but not homoneuronal, synaptic depression requires endocannabinoid signaling in astrocytes

We then investigated the role of astrocyte CB_1_Rs on the eCB-induced homoneuronal and heteroneuronal synaptic depression. We selectively deleted CB_1_R expression in cortical astrocytes by expressing Cre-recombinase under the control of the astroglial promoter GFAP, using local injection of AAV8-GFAP-mCherry-Cre in S1 of CB_1_R^flox/flox^ mice (**Figure 2A**). These mice are herein termed aCB_1_R^-/-^ mice, and their controls, termed aCB_1_R mice, were CB_1_R^flox/flox^ mice injected with AAV8-GFAP-mCherry (i.e., lacking Cre). To assess the efficacy of the approach, we monitored the CB_1_R-mediated astrocyte Ca^2+^ responses to the CB_1_R agonist WIN 55,212-2 (300 μM) using two-photon microscopy and the genetically encoded calcium indicator GCaMP6f selectively expressed in astrocytes via injection of AAV5-gfaABC1D-cyto-GCaMP6f in S1 (**Figure 2D**). While the astrocyte Ca^2+^ activity, quantified from the Ca^2+^ event probability, was increased by local application of WIN 55,212-2 in control aCB_1_R mice (n = 190 astrocytes from 13 slices), the WIN-evoked responses were significantly reduced in aCB_1_R^-/-^ mice (n = 261 astrocytes from 14 slices; **Figure 2D-F**), confirming the suitability of the viral approach to delete CB_1_R signaling in astrocytes.

**Figure 2.**
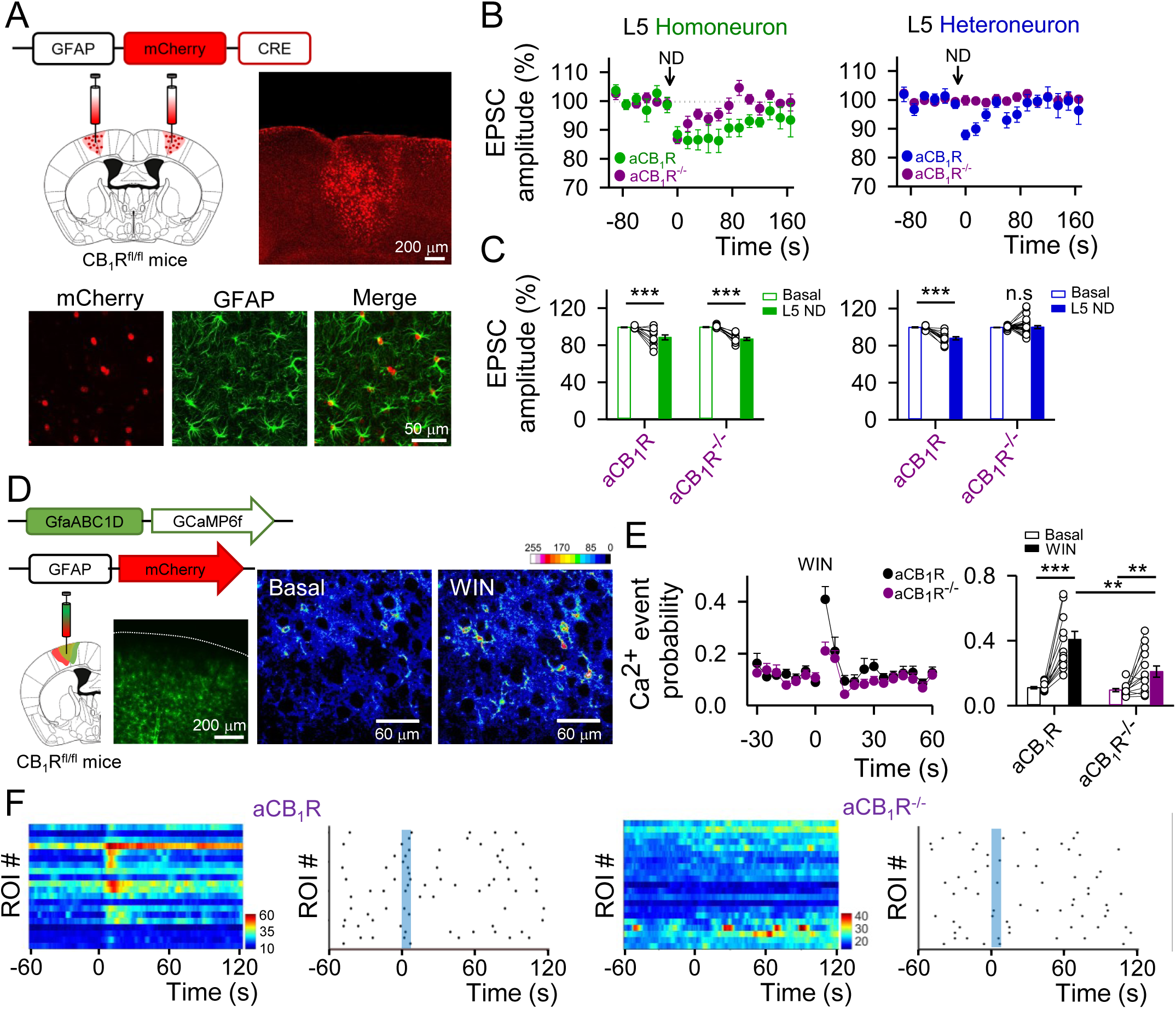
Heteroneuronal, but not homoneuronal, synaptic depression requires endocannabinoid signaling in astrocytes. (**A**) Viral vector injected into the S1 of CB_1_R^fl/fl^ mice and fluorescence image showing mCherry-Cre expression in the S1 (top), and immunohistochemistry images showing co-localization between mCherry-cre and GFAP (bottom). (**B**) EPSCs amplitude *versus* time before (basal) and after ND in CB_1_R mice injected with AAV8-GFAP-mCherry (aCB_1_R; green or blue) or with AAV8-GFAP-mCherry-Cre (aCB_1_R^-/-^; purple) in the homoneuron (left) and heteroneuron (right) from L5. (**C**) Relative changes in EPSC amplitude in aCB_1_R and aCB_1_R^-/-^ mice in the homoneuron (left) and heteroneuron (right). Two-tailed Student’s paired t test. (**D**) Viral vector injected into the S1 of CB_1_R^fl/fl^ mice, fluorescence image showing GCaMP6f expression in the S1 and pseudocolor images showing the fluorescence intensities of GCaMP6f-expressing astrocytes before and after WIN (300 μM) application in L5. (**E**) Ca^2+^ event probability over time (left) and Ca^2+^ event probability before (basal) and after WIN application in aCB_1_R (black) and aCB_1_R^-/-^ (purple) mice (right). Blue shadow indicates 5s WIN application. Two-tailed Student’s paired t test (before and after) and two-tailed Student’s unpaired t test (between groups). (**F**) Raster plots and heat maps showing the Ca^2+^ events recorded from all ROIs including astrocyte somas and processes in aCB_1_R (left) and aCB_1_R^-/-^ (right) mice before and after WIN stimulation. Blue shadow indicates 5s WIN application. Data are expressed as mean ± SEM, **p < 0.01, ***p < 0.001.

Next, we tested the impact of astroglial deletion of CB_1_Rs on the ND-evoked homoneuronal and heteoneuronal synaptic depression in L5. Accordingly, the homoneuronal depression was not affected in mice lacking CB_1_Rs in astrocytes (12 out of 35 cells; 34.3%) (**Figures 2B and 2C**). By contrast, the heteroneuronal synaptic depression was absent in aCB_1_R^-/-^ mice (0 out of 27 cells; 0%) (**Figures 2B and 2C**). These results indicate that eCB-induced heteroneuronal, but not homoneuronal, synaptic depression involves CB_1_R signaling in astrocytes.

### Heteroneuronal synaptic depression requires astrocyte Ca^2+^ signaling and activation of presynaptic A1 receptors

Ca^2+^ elevations in astrocytes evoked by different neurotransmitters, including eCBs^22,35,36^, are known to stimulate the release of gliotransmitters that regulate synaptic function (e.g.,^15,22,35–37^). Hence, we investigated whether the homoneuronal and heteroneuronal synaptic depressions depended on the astrocytic Ca^2+^ signal. We depolarized L5 pyramidal neurons using the approach that elicits homoneuronal and heteroneuronal synaptic depression as indicated above and monitored the astrocyte Ca^2+^ activity using GCaMP6f selectively expressed in astrocytes (**Figure 2D**). To isolate eCB-induced effects, experiments were performed in the presence of a cocktail of antagonists of glutamatergic, GABAergic, purinergic, and cholinergic receptors (see Material and Methods). Under these conditions, ND elevated astrocyte Ca^2+^ fluctuations (n = 142 astrocytes of 6 slices), an effect that was abolished in the presence of AM251, indicating that these responses were mediated by CB_1_R activation (n = 122 astrocytes of 6 slices; **Figures 3A and 3B**).

**Figure 3.**
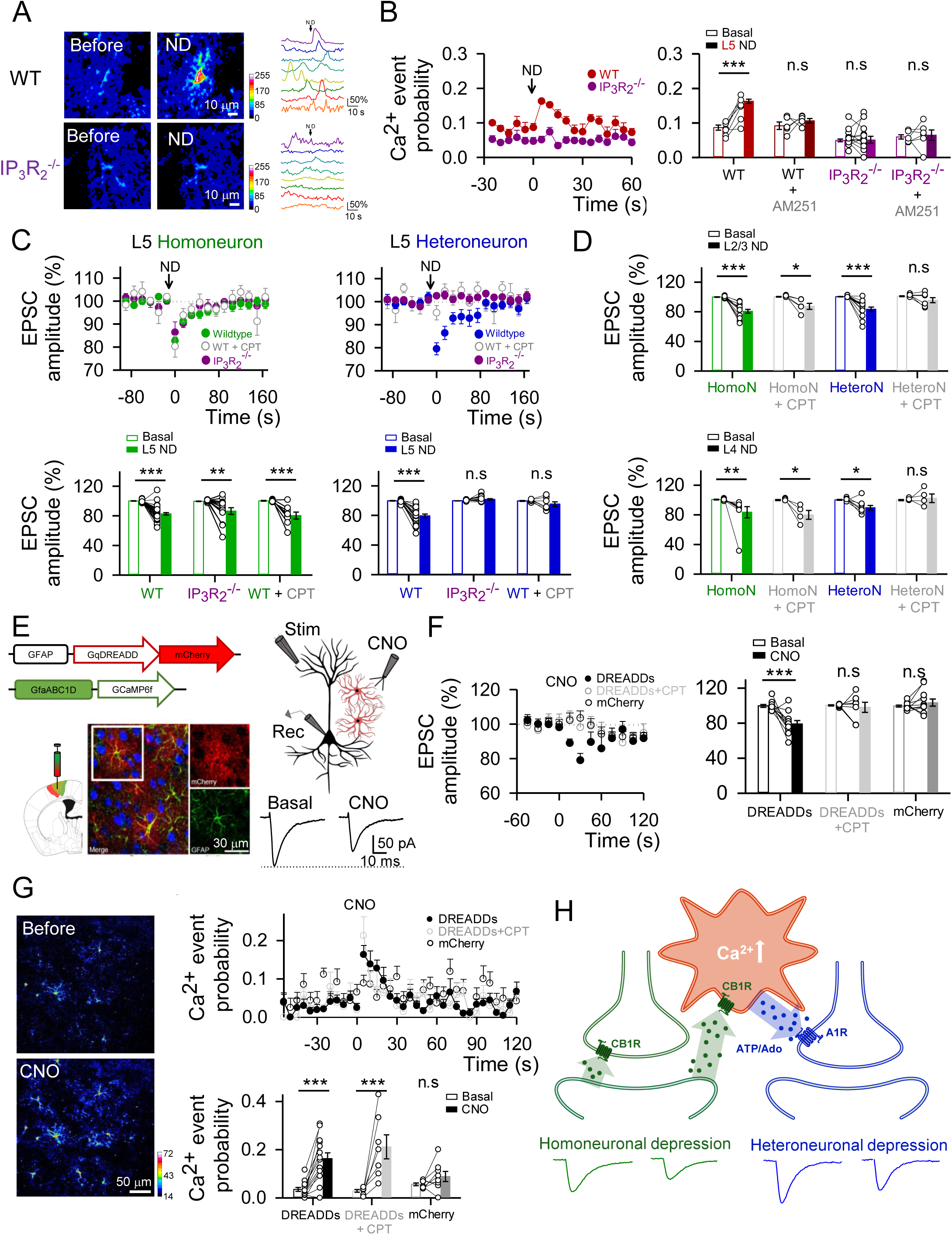
Heteroneuronal synaptic depression requires astrocyte calcium signaling and activation of presynaptic A1 receptors. (**A**) Left: Pseudocolor images showing the fluorescence intensities of GCaMP6f-expressing astrocytes in L5 before and after L5 ND in wildtype (top) and IP_3_R_2_^-/-^ mice (bottom). Right: representative Ca^2+^ traces of astrocytes (arrow indicates ND). (**B**) Left: L5 astrocytes Ca^2+^ event probability over time before (basal) and after L5 ND in wildtype (red) and IP_3_R_2_^-/-^ (purple) mice. Right: Relative changes in Ca^2+^ event probability in wildtype and IP_3_R_2_^-/-^ mice in control and with AM251 (2 μM). All experimental conditions were performed in TTX (1 μM) and in a cocktail of neurotransmitter receptor antagonists (see Material and Methods). Two-tailed Student’s paired t test. (**C**) Top: EPSCs amplitude *versus* time before (basal) and after ND in wildtype mice in control (green or blue), in presence of CPT (5 μM) (open gray) and in IP_3_R_2_^-/-^ mice (purple) in the homoneuron (left) and heteroneuron (right) from layer 5. Bottom: Relative changes in EPSC amplitude in wildtype mice in control, in presence of CPT (5 µM) and in IP_3_R_2_^-/-^ mice. Two-tailed Student’s paired t-test. (**D**) Top: Relative changes in EPSC amplitude before (basal) and after L2/3 ND in control and with CPT (5 µM) in the homoneuron (green) and heteroneuron (blue) from layer 2/3. Bottom: Relative changes in EPSC amplitude before (basal) and after L4 ND in control and with CPT (5 µM) in the homoneuron (green) and heteroneuron (blue) from layer 4. Two-tailed Student’s paired t test. (**E**) Left: Viral vectors injected into the S1 of wildtype mice and immunohistochemistry images showing the expression of NeuN (blue), mCherry (red) and GFAP (green) in the somatosensory cortex slices of a DREADDs injected mouse. Note the selective expression of hM3D–mCherry in astrocytes. Right: Scheme of the experimental approach and representative EPSC traces before (basal) and after CNO (1mM) application in L5. (**F**) Left: EPSCs amplitude *versus* time before (basal) and after CNO application in AAV8-GFAP-Gq-DREADD-mCherry injected mice in control (black, close) and in presence of CPT (gray, open), and in AAV8-GFAP-mCherry injected mice (black, open). Blue shadow indicates 5s CNO application. Right: Relative changes in EPSC amplitude in DREADDs injected mice in control and in presence of CPT, and in mCherry injected mice. Two-tailed Student’s paired t test. (**G**) Left: Pseudocolor images showing the fluorescence intensities of GCaMP6f-expressing astrocytes before and after CNO application in L5. Top right: Ca^2+^ event probability over time of L5 astrocytes before (basal) and after CNO application in AAV8-GFAP-Gq-DREADD-mCherry injected mice in control (black, close) and in presence of CPT (gray, open), and in AAV8-GFAP-mCherry injected mice (black, open). Blue shadow indicates 5s CNO application. Bottom right: relative changes in Ca^2+^ event probability in DREADDs injected mice in control and in presence of CPT, and in mCherry injected mice. Two-tailed Student’s paired t test. (**H**) Schematic summary depicting the signaling pathways involved in eCBs-induced heteroneuronal synaptic depression. Data are expressed as mean ± SEM, *p < 0.05, **p < 0.01, ***p < 0.001.

Moreover, ND-evoked astrocyte Ca^2+^ elevations were largely absent in inositol-1,4,5-trisphosphate (IP3)-receptor type 2-deficient mice (IP_3_R_2_^-/-^ mice), in which G protein-mediated Ca^2+^ signal is selectively impaired in astrocytes^27,38–40^ (n = 164 astrocytes of 11 slices; **Figures 3A and 3B**). In these mice, the homoneuronal depression was preserved (13 out of 30 cases; 43.3%), but the heteroneuronal depression was absent (0 out of 17 cases; 0%) (**Figure 3C**). Collectively these results indicate that the astrocyte Ca^2+^ signal is required for the heteroneuronal, but not the homoneuronal, synaptic depression.

We then investigated the gliotransmitter responsible for the heteroneuronal depression. ATP and its metabolic product adenosine are known to be released by astrocytes^8^ and to regulate synaptic transmission in several brain areas^23,24,36,37,41,42^. Therefore, we hypothesized that eCB-induced astrocyte calcium elevations would stimulate the release of ATP/adenosine that acting on neuronal type 1 adenosine receptors (A1Rs) would depress synaptic transmission. To test this idea, we depolarized L5 pyramidal neurons and monitored the homo- and heteroneuronal synaptic depression before and after bath application of the A1R antagonist CPT (5 μM). While the homoneuronal depression was unaffected (n = 8), the heteroneuronal depression was abolished in the presence of CPT (n = 8; **Figure 3C**). Consistent with results observed in L5, heteroneuronal depressions in L4 and L2/3 were also abolished by the A1R antagonist CPT (**Figure 3D**).

To further test the astrocyte involvement, we investigated if activation of G-protein-mediated signaling in astrocytes depresses excitatory transmission in the S1 cortex by directly and selectively activating astrocytes using designer receptors exclusively activated by designed drugs (DREADDs). Astrocytes in the S1 cortex were targeted with AAV8-GFAP-Gq-DREADD-mCherry and AAV5-gfaABC1d-GCaMP6f (**Figure 3E**). Activation of Gq-DREADDs with the synthetic agonist clozapine-N-oxide (CNO, 1 mM) delivered from a micropipette by pressure pulses (5 s) (**Figure 3E**) induced Ca^2+^ elevations in astrocytes (n = 253 astrocytes from 14 slices, **Figure 3G**) and depressed synaptic transmission in L5 pyramidal neurons (n = 11; **Figure 3F**), which was associated with an increase in PPR indicating a presynaptic mechanism (n = 11; **Figure S1E**). Moreover, in the presence of CPT, CNO also induced Ca^2+^ elevations in astrocytes (n = 87 astrocytes from 7 slices, **Figure 3G**) but failed to affect synaptic transmission (n = 7, **Figure 3F**). In slices from mice that were injected with control AAV8-GFAP-mCherry virus (i.e., lacking DREADDs), CNO application failed to affect both synaptic transmission (n = 11; **Figure 3F**) and astrocyte Ca^2+^ dynamics (n = 92 astrocytes from 8 slices; **Figure 3G**). These results suggest that astrocyte Ca^2+^ elevations are sufficient to regulate cortical synaptic transmission.

Taking together, these results indicate that neuron-released eCBs induce homoneuronal depression by directly acting on neuronal CB_1_Rs. Concomitantly, eCBs activate CB_1_Rs in astrocytes, elevate their intracellular Ca^2+^, and stimulate the release of ATP/Adenosine, which acts on presynaptic A1Rs triggering the heteroneuronal depression (**Figure 3H**).

### Astrocyte-mediated heteroneuronal depression is column-specific

The functional properties of the somatosensory cortex rely on their modular organization, comprising sub-circuits of layer connectivity within cortical columns^3,5,6^. Thus, we investigated whether the astrocyte-mediate heteroneuronal depression was also spatially restricted to a single column. We analyzed this phenomenon in the somatosensory barrel cortex, where cortical columns can be easily identified (**Figure 4A**). We performed paired whole-cell recordings of L4 neurons located at a similar distance (70-270 μm) but either within the same or in adjacent columns (**Figure 4A**). Depolarization of a single L4 neuron to stimulate eCBs release induced heteroneuronal depression in the paired recorded neuron located in the same column (10 out of 29 paired recordings; 35%, **Figure 4B**). Consistent with the mechanistic interpretation described above, this phenomenon was blocked after CPT perfusion (**Figure S1G**). In contrast to this intracolumn regulation, the heteroneuronal regulation was absent in neurons located at a similar distance but in an adjacent cortical column (0 out 15 paired recordings; 0%, **Figure 4C**). In both cases, intra- and intercolumn recordings displayed homoneuronal synaptic depression (16 out 48 cells; 33%, **Figure S1F**).

**Figure 4.**
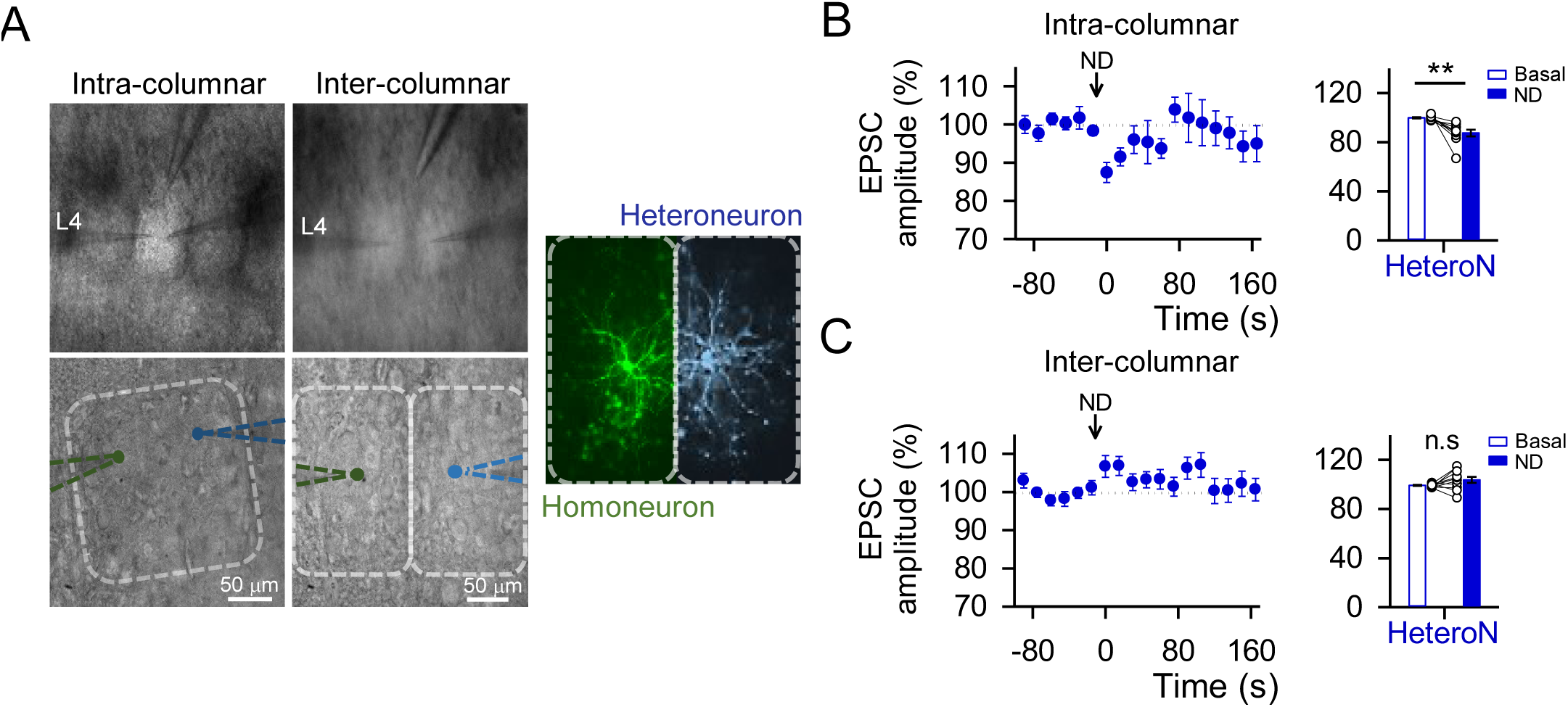
Astrocyte-mediated heterosynaptic depression is column specific. (**A**) Representative infrared differential interference contrast images of the barrel field in the primary somatosensory cortex showing intracolumn (left) and intercolumn (middle) pair of neurons patched and the stimulation electrode. False-color biocytin loading barrel cortex intercolumnar pair of neurons image (right). (**B**) Left: EPSCs amplitude *versus* time before (basal) and after ND in the intracolumn heteroneuron. Right: Relative changes in EPSC amplitude in the intracolumn heteroneuron. (**C**) Left: EPSCs amplitude *versus* time before (basal) and after ND in the intercolumn heteroneuron. Right: Relative changes in EPSC amplitude in the intercolumn heteroneuron. Two-tailed Student’s paired t test. Data are expressed as mean ± SEM, **p < 0.01.

Overall, these results indicate that the eCB-induced astrocyte-mediated heteroneuronal synaptic regulation is column-specific, i.e., it is not a wide unspecific phenomenon but a synaptic regulatory signaling that specifically occurs between cells located within a cortical column.

### Astrocyte-mediated heterosynaptic depression is layer-specific

Cortical information processing depends not only on the columnar organization but also on the functional interaction across different layers^6^. Therefore, we examined the functional organization of heteroneuronal and homoneuronal synaptic depression across different cortical layers, i.e., between neurons located in L2/3, L4, and L5 (**Figure 5**). We performed paired recordings of neurons in these layers, depolarized one neuron to stimulate eCB release, and monitored EPSCs in the other neuron located in another layer. While depolarizing a single L2/3 neuron did not affect synaptic transmission in L4 neurons (n = 11) (**Figures 5A and 5B**), stimulation of a single L4 neuron induced heteroneuronal synaptic depression in L2/3 neurons (5 out of 10 cases; 50%, **Figures 5A and B**). Likewise, stimulation of a single L2/3 neuron did not alter synaptic transmission in L5 neurons (n = 8; **Figures 5C and 5D**), but stimulation of L5 neurons induced heteroneuronal depression in L2/3 neurons (4 out of 10 cases; 40%, **Figures 5C and 5D**). Finally, stimulation of L4 neurons depressed neurotransmission in L5 pyramidal neurons (4 out of 14 cases; 28.1%, **Figures 5E and 5F**), but L5 neuron stimulation did not impact synaptic transmission in L4 neurons (n = 16; **Figures 5E and 5F**). In summary, astrocyte-mediated heteroneuronal depression occurs between neurons located in different layers, but following a specific pattern and not necessarily reciprocally. For example, synapses in L2/3 neurons can be regulated by neurons located in L4 or L5, but not vice versa; and L4 neurons can regulate neurons in L2/3 and L5 but are not regulated by them.

**Figure 5.**
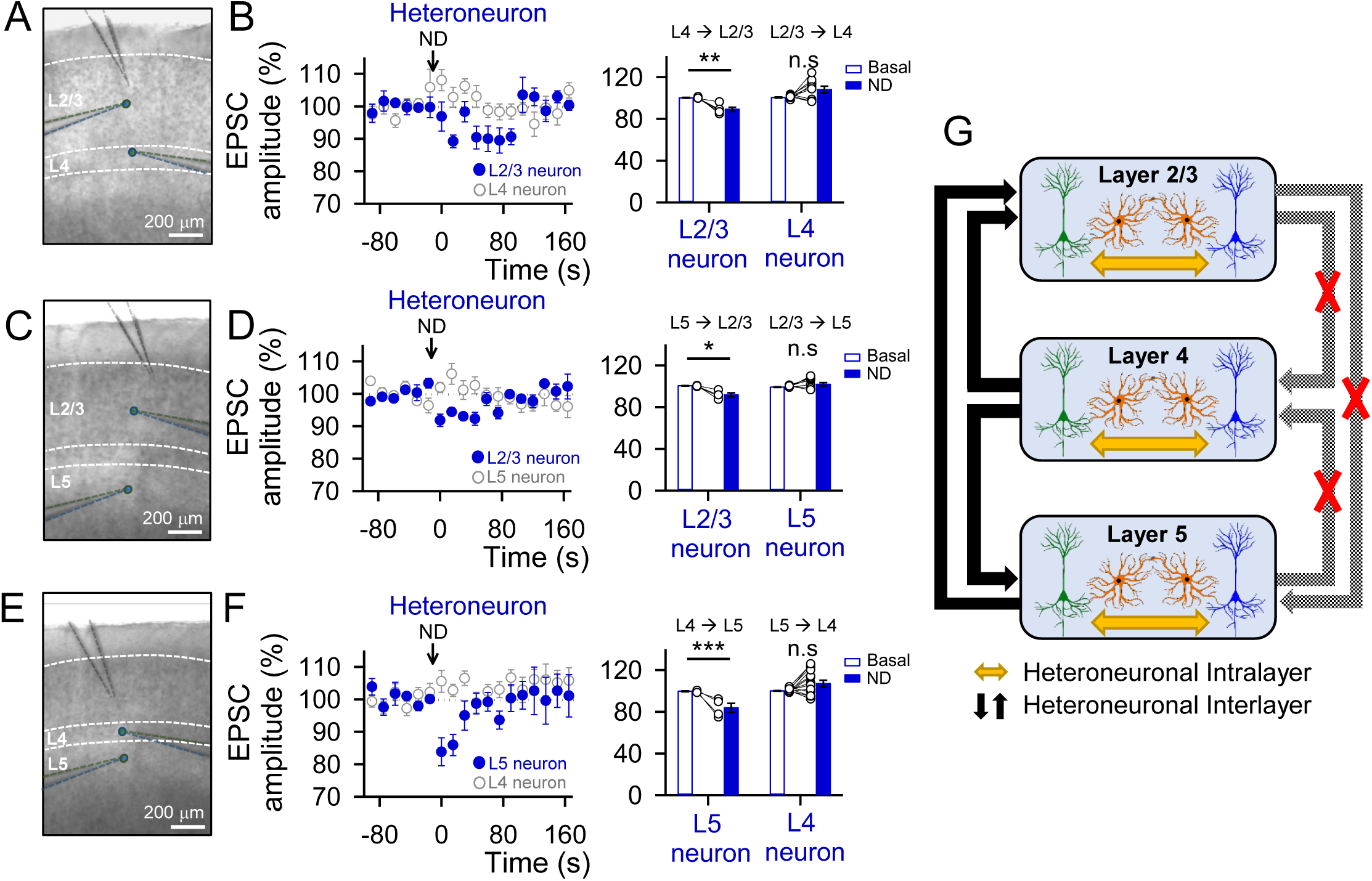
Astrocyte-mediated heterosynaptic depression is layer specific. (**A**) Representative infrared differential interference contrast image of the experimental configuration with the stimulation pipette in layer 2/3 and a pair of neurons patched one in L4 and the other in L2/3. (**B**) Left: Heteroneuronal EPSCs amplitude *versus* time in a pair of neurons patched one in L4 and the other in L2/3 before (basal) and after ND of L4 (blue) or L2/3 neuron (open grey). Right: Relative changes in EPSC amplitude in L2/3 (when L4 neuron is stimulated) and L4 (when L2/3 neuron is stimulated) neuron. Two-tailed Student’s paired t-test. (**C**) Representative infrared differential interference contrast image of the experimental configuration with the stimulation pipette in layer 2/3 and a pair of neurons patched one in L5 and the other in L2/3. (**D**) Left: Heteroneuronal EPSCs amplitude *versus* time in a pair of neurons patched one in L5 and the other in L2/3 before (basal) and after ND of L5 (blue) or L2/3 neuron (open grey). Right: Relative changes in EPSC amplitude in L2/3 (when L5 neuron is stimulated) and L5 (when L2/3 neuron is stimulated) neuron. Two-tailed Student’s paired t-test. (**E**) Representative infrared differential interference contrast image of the experimental configuration with the stimulation pipette in layer 2/3 and a pair of neurons patched one in L4 and the other in L5. (**F**) Left: Heteroneuronal EPSCs amplitude *versus* time in a pair of neurons patched one in L4 and the other in L5 before (basal) and after ND of L4 (blue) or L5 neuron (open grey). Right: Relative changes in EPSC amplitude in L5 (when L4 neuron is stimulated) and L4 (when L5 neuron is stimulated) neuron. Two-tailed Student’s paired t test. (**G**) Schematic summary depicting the astrocyte-mediated heterosynaptic regulation pathways into the same layer (intralayer) and between layers (interlayer). Data are expressed as mean ± SEM, *p < 0.05, **p < 0.01, ***p < 0.001.

Together, these results indicate that eCB-induced astrocyte-mediated heteroneuronal synaptic regulation is not an unspecific phenomenon, rather it is layer-specific, selectively occurring among neurons following a layer-specific pattern (**Figure 5G**).

### Astrocytic calcium responses to eCBs are not homogeneous across cortical layers

Because the heteroneuronal depression depends on the eCB-induced astrocyte Ca^2+^ signals, its layer-specificity might be accounted for by layer-specificity of astrocyte responsiveness to eCBs (**Figure 6A**). To test this idea, we examined the astrocyte Ca^2+^ signals in different layers in response to eCBs released by depolarization of neurons. To ensure that the astrocyte activation was due to eCBs, we performed the experiments in the presence of TTX (1 μM) and the cocktail of neurotransmitter receptor antagonists (see Material and Methods). Neuronal depolarization of L2/3, L4 or L5 neurons elevated intracellular Ca^2+^ in astrocytes located within the same layer (n = 110 astrocytes from 5 slices; n = 110 astrocytes from 4 slices; n = 142 from 6 slices respectively). These responses were abolished by AM251, confirming to be the result of eCB signaling (n = 89 astrocytes from 5 slices; n = 79 astrocytes from 4 slices; n = 122 from 6 slices respectively, **Figures 6B-D**).

**Figure 6.**
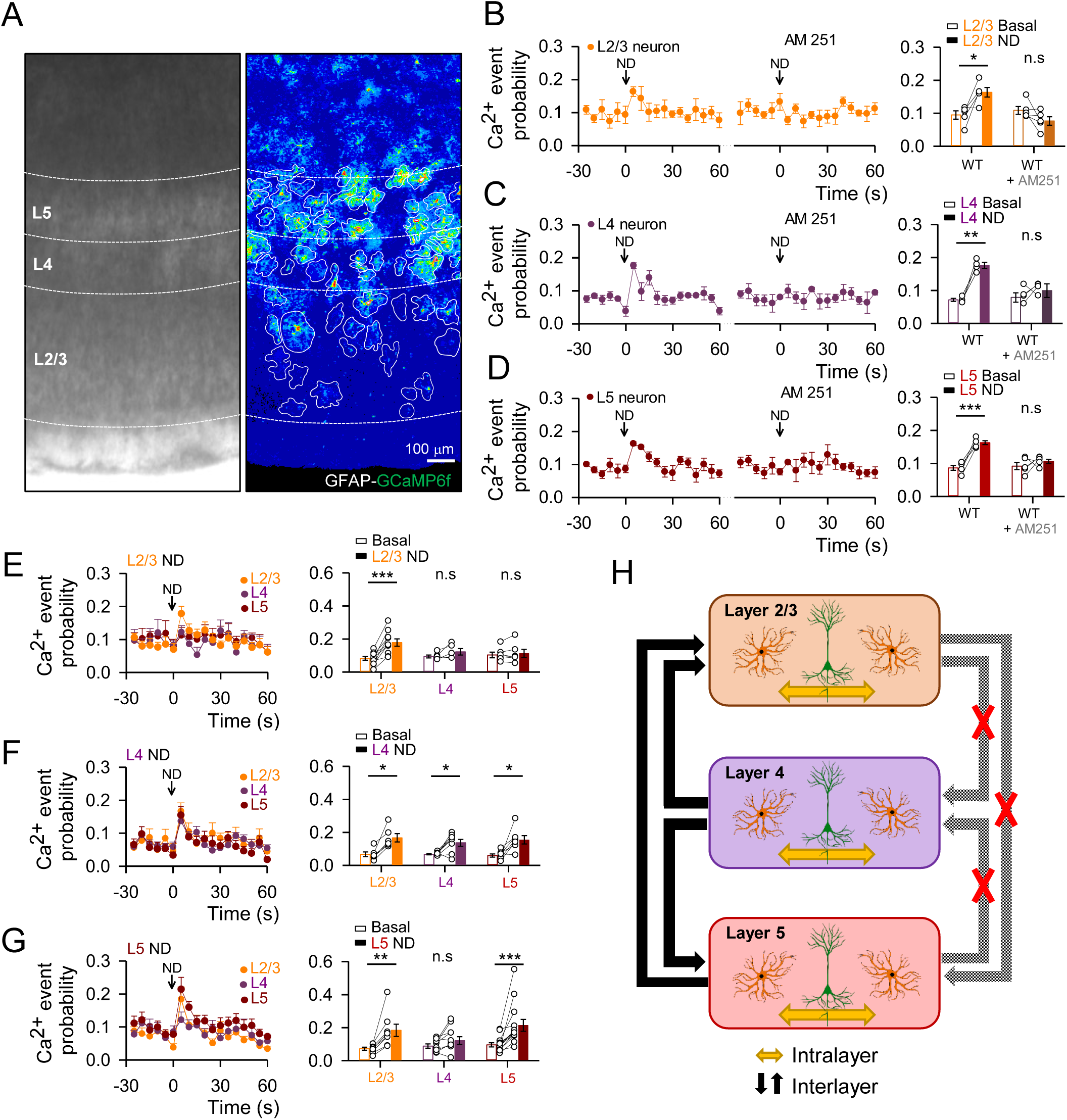
Astrocytic calcium responses to eCBs are not homogeneous across cortical layers. (**A**) Representative infrared differential interference contrast image and pseudocolor image representing fluorescence intensities of GCaMP6f-expressing astrocytes in the different layers of the primary somatosensory cortex. (**B-D**) Left: Ca^2+^ event probability over time before (basal) and after ND in control and in presence of AM251 (2 μM) when patched and recorded in L2/3 (**B**, orange), L4 (**C**, purple) or L5 (**D**, red). Right: relative changes in Ca^2+^ event probability in control and in presence of AM251 (2 μM) when patched and recorded in L2/3 (**B**, orange), L4 (**C**, purple) or L5 (**D**, red). Two-tailed Student’s paired t test. (**E**) Left: Ca^2+^ event probability over time of astrocytes of layer 2/3 (orange), 4 (purple) and 5 (red) before (basal) and after L2/3 neuron depolarization. Right: relative changes in Ca^2+^ event probability of astrocytes of layer 2/3 (orange), 4 (purple) and 5 (red). Two-tailed Student’s paired t test. (**F**) Left: Ca^2+^ event probability over time of astrocytes of layer 2/3 (orange), 4 (purple) and 5 (red) before (basal) and after L4 neuron depolarization. Right: relative changes in Ca^2+^ event probability of astrocytes of layer 2/3 (orange), 4 (purple) and 5 (red). Two-tailed Student’s paired t test. (**G**) Left: Ca^2+^ event probability over time of astrocytes of layer 2/3 (orange), 4 (purple) and 5 (red) before (basal) and after L5 neuron depolarization. Right: relative changes in Ca^2+^ event probability of astrocytes of layer 2/3 (orange), 4 (purple) and 5 (red). Two-tailed Student’s paired t test. All experimental conditions were performed in TTX (1 μM) and in a cocktail of neurotransmitter receptor antagonists (see Material and Methods). (**H**) Schematic summary depicting the calcium responses of astrocytes located in the same (intralayer) or different (interlayer) layers to the endogenous mobilized eCBs from neurons located in the same or different layers. Data are expressed as mean ± SEM, *p < 0.05, **p < 0.01, ***p < 0.001.

We then examined the astrocyte responses to neuron-released eCBs across cortical layers. Stimulation of L2/3 neurons increased the astrocyte Ca^2+^ event probability in L2/3 (n = 185 astrocytes from 10 slices) but failed to increase Ca^2+^ signaling in L4 (n = 62 astrocytes from 5 slices) or L5 (n = 129 astrocytes from 6 slices) astrocytes (**Figure 6E**). Likewise, stimulation of L4 neurons elevated Ca^2+^ in L4 (n = 163 astrocytes from 8 slices), L5 (n = 87 astrocytes from 7 slices) and L2/3 (n = 71 astrocytes from 6 slices) astrocytes (**Figure 6F**). Finally, stimulation of L5 neurons produced an increase in the Ca^2+^ event probability in L5 (n = 267 astrocytes from 13 slices) and L2/3 (n = 202 astrocytes from 8 slices) astrocytes but not in L4 (n = 144 astrocytes from 8 slices) astrocytes (**Figure 6G**). These results indicate that astrocyte Ca^2+^ increases mediated by eCBs signaling obeys a layer-specific pattern in agreement with the astrocyte-mediated heterosynaptic regulation produced by neuronal depolarization (**Figure 6H**).

In summary, eCB-mediated neuron-to-astrocyte signaling is a form of communication that occurs between cells that can be located in different layers, but following a specific connectivity pattern. Like the heteroneuronal depression, this specific pattern does not necessarily involve reciprocal signaling between layers. Notably, the specific neuron-to-astrocyte connectivity pattern mirrors the heteroneuronal depression pattern, suggesting that the selective neuron-astrocyte signaling between layers is responsible for the astrocyte-mediated non-synaptic communication between neurons in different cortical layers.

## DISCUSSION

Present results show that astrocytes regulate cortical synaptic function in a layer- and column-specific manner and that the functional interaction between cortical astrocytes and synapses is highly spatially organized. We show that activation of astrocytes by endogenous stimuli –eCBs physiologically released from cortical neurons–, induced astrocyte Ca^2+^ elevations and transiently depressed synaptic transmission in neurons located in the same and distinct cortical layers. This heteroneuronal synaptic depression requires astrocytic CB_1_R activation, is mediated by activation of presynaptic A1 receptors, and can be mimicked by astrocyte-specific chemogenetic stimulation. Additionally, our results also show that neuron-released eCBs can depress synaptic transmission by directly activating CB_1_Rs in homoneuronal synapses, a phenomenon known as DSE^29–34^.

The neocortex is highly organized in layers and columns with precisely neuronal connectivity. Our results indicate that eCB-mediated astrocyte-neuron signaling is also exquisitely organized. First, the astrocyte-mediated heteroneuronal depression was found to be column-specific because it only occurred between neurons located within the same column and not between neurons located at similar distances but in adjacent columns (**Figure 4**). Second, astrocyte-mediated heteroneuronal regulation occurred between neurons located in different layers, but according to a specific connectivity pattern (**Figure 5**). Third, the eCB-mediated neuron-to-astrocyte signaling was also layer-specific because astrocytic calcium responses to eCBs released by neurons in different cortical layers were not homogeneous across the cortical layers; rather neuron-to-astrocyte signaling occurred according to particular signaling patterns (**Figure 6**).

Several previous studies have shown that astrocytes stimulated by eCBs lead to regulation of synaptic transmission in diverse brain areas, including the hippocampus, amygdala, and neocortex^15,22,26,27^. In the neocortex, synapses onto layer 2/3 neurons undergo spike-timing long-term depression (LTD) mediated by glutamate released from astrocytes^15^. By contrast, we found (**Figures 1, 2, and 3**) that eCB-induced astrocyte activation transiently depresses synapses through ATP/adenosine release as gliotransmitters. Different neuronal stimulating paradigms used in these studies may account for such discrepancies. As a matter of fact, astrocytes are competent to release distinct gliotransmitters depending on the pattern of neuronal stimulation as demonstrated in the hippocampus, where astrocytes can release glutamate upon low frequency stimulation of neighboring interneurons or glutamate and ATP/adenosine upon high frequency stimulation^43^.

The eCB-induced astrocyte-mediated heteroneuronal depression was found to be restricted within a single cortical column, supporting the idea of a highly organized signaling between astrocytes and neurons at a modular level. These results agree with previous reports showing that astrocyte Ca^2+^ signal is spatially restricted in astrocytes located within the columns of the barrel cortex^14,44,45^. The column-specific astrocyte-mediated synaptic regulation also indicates that astrocyte-neuron networks are functionally organized obeying the columnar organization of the neuronal connectivity pattern.

In conclusion, the present data indicate that astrocytes modulate cortical synaptic transmission in a column and layer-specific manner, obeying the structural and functional organization of the cortex, which suggests that they are an integral part of the cortical modules. Moreover, astrocytes, by providing layer-specific signaling pathways of non-synaptic communication between neurons, add further complexity to the signaling mechanisms underlying cortical network function. This finely controlled astrocyte-synapse interaction is particularly significant in the neocortex, where the spatial integration of synaptic signals is highly relevant for cortical information processing.

## METHODS

### Ethics statement

All of the procedures for handling and sacrificing animals were approved by the University of Minnesota Institutional Animal Care and Use Committee (IACUC) in compliance with the National Institutes of Health guidelines for the care and use of laboratory animals.

### Animals

Mice were housed under 12/12-h light/dark cycle, up to five animals per cage, at temperatures between 68–74°F at 30–70% humidity with freely available food and water. The following animals (males and females) were used for the present study C57BL/6J, IP_3_R_2_^-/-^ (generously donated by Dr. J Chen), and CB_1_R^fl/fl 46,47^. Adult (≥8 weeks) mice were used.

### Somatosensory Cortex Slice Preparation

Mice were euthanized by decapitation and brains were rapidly removed and placed in ice-cold artificial cerebrospinal fluid (ACSF). Three-hundred and fifty-micrometer coronal brain slices containing the somatosensory cortex were prepared via a Leica VT1200 vibratome in a 4°C ACSF solution. Following cutting, slices were allowed to recover in ACSF containing (in mM): NaCl 124, KCl 2.69, KH_2_PO_4_ 1.25, MgSO_4_ 2, NaHCO_3_ 26, CaCl_2_ 2 and glucose 10, gassed with 95% O_2_/5% CO_2_ (pH = 7.3) at 31°C for 30 min followed by 30 min at 20–22°C before recording. After a 1 h recovery period, slices were kept at 20–22°C for the rest of the recording day. Slices were then transferred to an immersion recording chamber and superfused at 2 ml/min with gassed ACSF and the temperature of the bath solution was kept at 34ºC with a temperature controller TC-324B (Warner Instruments Co.). Cells were visualized using infrared-differential interference contrast optics (Nikon Eclipse E600FN, Tokyo, Japan) and 40x water immersion lens. L2/3, L4, and L5 from the forelimb and hindlimb somatosensory cortex and the barrel subfields were identified with a 10x objective.

### Electrophysiology

Neurons were selected based on their location, morphology, and firing pattern. Simultaneous dual electrophysiological recordings from layers 2/3, 4, and 5 pyramidal neurons were made using the whole-cell-patch-clamp technique. When filled with an internal solution containing (in mM): KGluconate 135, KCl 10, HEPES 10, MgCl_2_ 1, ATP-Na_2_ 2 (pH = 7.3) patch electrodes exhibited a resistance of 3-10 MΩ. All recordings were performed using PC-ONE amplifiers (Dagan Instruments, Minneapolis, MN). Fast and slow whole-cell capacitances were neutralized, and series resistance was compensated (∼70%), and the membrane potential was held at -70 mV. Intrinsic electrophysiological properties were monitored at the beginning and the end of the experiments. Series and input resistances were monitored throughout the experiment using a -5 mV pulse. Recordings were considered stable when the series and input resistances, resting membrane, and stimulus artifact duration did not change > 20%. Furthermore, I-V curves and firing patterns at the beginning and the end of the experiments were similar. Recordings that did not meet these criteria were discarded. Signals were fed to a Pentium-based PC through a DigiData 1322A interface board. Signals were filtered at 1 kHz and acquired at a 10 kHz sampling rate using a DigiData 1322A data acquisition system and pCLAMP 10.3 software (Molecular Devices, San Jose, CA). Distance of the somas of the paired recorded neurons within a layer varied from, 70-270 µm. In paired recordings across layers 2/3, 4, and 5, neurons were selected following the same vertical axis.

### Synaptic Stimulation

Theta capillaries (2-5 µm tip) filled with ACSF were used for bipolar stimulation. The electrodes were connected to a stimulator S-900 through an isolation unit and placed in L2/3. When indicated, the stimulation electrode was placed in L4. Paired pulses of 1 ms duration and 50 ms interval were continuously delivered at 0.33 Hz. Excitatory postsynaptic currents (EPSCs) were isolated using picrotoxin (50 μM) and CGP5462 (1 μM) to block GABA_A_R and GABA_B_R, respectively. EPSC amplitude was determined as the peak current amplitude (2–20 ms after stimulus) minus the mean baseline current (10-30 ms before stimulus). The paired-pulse ratio (PPR) was estimated as PPR = (2^nd^ EPSC/1^st^ EPSC).

To induce eCB release, pyramidal neurons were depolarized from −70 mV to 0 mV for 5 s (ND)^28,29^. Synaptic parameters were determined from 60 stimuli before (basal) and following ND. Baseline mean EPSC amplitude was obtained by averaging mean values obtained within 3 min of baseline recordings and mean EPSC amplitudes were normalized to baseline. The ND was applied 2.5 s after the last basal delivered pulse, and no pulses were presented during the ND. Immediately after the ND was finished, the 0.33-Hz pulse protocol was restarted. To illustrate the time course of ND-induced effects, synaptic parameters were grouped in 15 s bins. Three consecutive responses to ND were averaged. A response was considered a depression if the amplitude of the current was < 2 times the standard deviation of the baseline current during the first 45 s after ND and was verified by visual inspection.

The effects of pharmacological agents (CPT 5 μM and AM251 2 μM) were tested after 10 min bath perfusion and in the same neurons that previously were depolarized in control conditions. In all cases the effects of pharmacological agents were tested at < 40 min after entering whole-cell mode in the stimulating neuron.

### Ex vivo two-photon calcium fluorescence imaging and electrophysiology

Two-photon microscopy imaging was performed using a Leica SP5 multi-photon microscope (Leica Microsystems, USA) controlled by the Leica LAS software and adapted to perform electrophysiological recordings. C57BL/6J, IP_3_R_2_^-/-^ and aCB_1_R mice injected into S1 with AAV5-GfaABC1d-GCaMP6f and AAV8-GFAP-mCherry were used (for aCB_1_R^-/-^ instead AAV8-GFAP-mCherry we used AAV8-GFAP-mCherry-Cre). All Ca^2+^ experiments, except those in which synaptic transmission was recorded, were performed in the presence of TTX (1 μM) and a cocktail of neurotransmitter receptor antagonists containing: CNQX (20 μM), AP5 (50 μM), MPEP (50 μM), LY367385 (100 μM), picrotoxin (50 μM), CGP5462 (1 μM), atropine (50 μM), CPT (5 μM), flupenthixol (30 μM), and suramin (100 μM).

Videos were obtained at 512 × 512 resolution with a sampling interval of 1 s. Red and green fluorescence was obtained in parallel to match red mCherry-stained astrocyte structure with green GCaMP6f astrocyte calcium. A custom MATLAB program (Calsee: https://www.araquelab.com/code/) was used to quantify fluorescence level measurements in astrocytes. Ca^2+^ variations recorded at the soma and processes of the cells were estimated as changes of the fluorescence signal over baseline (ΔF/F_0_), and cells were considered to show a Ca^2+^ event when the ΔF/F_0_ increase was at least two times the standard deviation of the baseline.

The astrocyte Ca^2+^ signal was quantified from the Ca^2+^ event probability, which was calculated from the number of Ca^2+^ elevations grouped in 5 s bins recorded from 8-50 astrocytes per field of view (layer 2/3, 4 or 5 of S1). The time of occurrence was considered at the onset of the Ca^2+^ event. For each astrocyte analyzed, values of 0 and 1 were assigned for bins showing either no response or a Ca^2+^ event, respectively, and the Ca^2+^ event probability was obtained by dividing the number of astrocytes showing an event at each time bin by the total number of monitored astrocytes^22^. All the astrocytes that showed a Ca^2+^ event during the experiment were used for the analysis. The calcium event probability was calculated in each slice, and for statistical analysis, the sample size corresponded to the number of slices as different slices were considered as independent variables. To examine the difference in Ca^2+^ event probability in distinct conditions, the basal Ca^2+^ event probability (mean of the 30 s before a stimulus) was averaged and compared to the average Ca^2+^ event probability (5 s after a stimulus). For ND experiments, each layer was recorded 1 minute before and after the ND. Three consecutive responses to ND were averaged in each layer. For WIN application, a micropipette was filled with 300 μM WIN solution and placed 100– 150 µm away from the tissue (layer 5), and a pressure pulse at 1 bar (PMI-100 DAGAN, Minneapolis, MN) was applied for 5 s. The absence of mechanical movement of the tissue was confirmed in every case. For acute application of CNO, a micropipette was filled with 1 mM CNO solution and placed 100–150 µm away from the recording neuron, and a pressure pulse was applied for 5 s. The absence of mechanical movement of the tissue was confirmed in every case. Stimulus effects on EPSCs were statistically tested comparing the normalized EPSCs recorded 1 min before and 30 s after the stimulus to assess changes in EPSC amplitude and PPR. Astrocytic Ca^2+^ events were recorded at the same time. The changes on the Ca^2+^ event probability after CNO application were statistically tested comparing the basal Ca^2+^ event probability 1 minute before and 5 s after the stimulus.

The effects of pharmacological agents (CPT 5 μM and AM251 2 μM) were tested after 10 min bath perfusion in the same region and same astrocytes recorded in control conditions. In the cases when Ca^2+^ imaging and electrophysiology were performed at same time the effects of pharmacological agents were tested at < 40 min after entering whole-cell mode in the stimulating or recorded neuron.

### AAV viral surgeries

Animals were anesthetized using a ketamine (10 mg/mL) xylazine (1 mg/mL) mixture and placed on a heating pad to maintain body temperature and faux tears were applied to the cornea. Animals (8 weeks of age) were placed in a stereotaxic apparatus and an incision was made down the midline of the scalp to expose the skull. A hole was drilled over the forelimb and hindlimb somatosensory cortex (S1: −0.4_a–p_, 1.9_m–l_), and a Hamilton syringe was lowered to (in mm from bregma: −0.7_d–v_) and viruses were injected bilaterally at 100 nL/min^48^. Mice were then sutured and left to heal for 2–3 weeks.

AAV5-pZac2.1-gfaABC1d-cyto-GCaMP6f (Addgene), AAV8-GFAP-hM3D(Gq)-mCherry (UMN vector core), AAV8-GFAP-mCherry (UMN vector core) and AAV8-GFAP-mCherry-Cre (UMN vector core) viral constructs were used. For CNO experiments, C57BL/6J mice were injected with AAV8-GFAP-hM3D(Gq)-mCherry virus. In control conditions, a virus of AAV8-GFAP-mCherry was injected instead. For CB_1_R^fl/fl^ mice experiments, AAV8-GFAP-mCherry-Cre was injected to delete CB_1_R from astrocytes (aCB_1_R^-/-^). AAV8-GFAP-mCherry was used as a control (aCB_1_R).

### Immunohistochemistry

The animals were anesthetized with Avertin (2,2,2 tribromoethanol, 240 mg/kg, i.p.) and intracardially perfused with ice cold phosphate buffered saline (PBS) and subsequently with 4% paraformaldehyde (PFA) in 0.1 M phosphate buffered saline (pH 7.4). The brain was removed, and 100 um coronal sections were made using a Leica VT1000S vibratome. Vibratome sections were incubated for one h in blocking buffer (0.1% Triton X-100, 10% Donkey or Goat serum in PBS) at room temperature. The primary antibodies were diluted in the blocking solution and the sections were incubated for two days at 4ºC. The following primary antibodies were used: Rabbit anti-GFAP (Sigma, 1:500) Mouse anti-NeuN (Millipore, 1:500). The slices were then washed three times for fifteen minutes each in PBS. The secondary antibodies were diluted in the secondary antibody buffer (0.1% Triton X-100, 5% Donkey or Goat serum in PBS) and incubated for 2 days at room temperature. The following secondary antibodies were used: 488 goat anti-rabbit (Invitrogen, 1:1000), 405 goat anti–mouse (Invitrogen, 1:500). The sections were then washed 3 times with 1xPBS for 10 min each and mounted using Vectashield Mounting media (Vector laboratories). The slides were imaged using a Leica SP5 multiphoton confocal microscope and Olympus FluoView FV1000.

The cellular specificity of Cre viral vectors was tested by immohistochemical analysis of randomly selected areas of the S1. Out of the 784 cells expressing mCherry from the AAV8-GFAP-mCherry-Cre viral vector, 86.7% were astrocytes (identified by GFAP) and 11.3% were neurons (identified by NeuN) (**Figures S2A and S2B**).

### Biocytin-stained neurons

Pair of neurons were recorded with patch pipettes and filled with internal solution containing 0.5% biocytin. Slices were fixed in 4% PFA in 0.1 PBS (pH 7.4) at 4ºC. Slices were washed three times in 1xPBS (10 min each). To visualize biocytin slices were incubated with Alexa488-Streptavidin (RRID: AB 2315383; 1:500) for 48 h at 4ºC. Slices were then washed for 3 times with 1xPBS (10 min each) and mounted with Vectashield mounting media (Vector laboratories). All mounted slices were imaged using a Leica SP5 multi-photon microscope. Also, pair of neurons were filled with biocytin through whole-cell recording, the slices were fixed using 4% paraformaldehyde. Then the slices were washed with PBS (100 mM sodium phosphate, pH 7.2). Endogenous peroxidases were then quenched by incubation with 1% H2O2. The slices were subsequently rinsed in PBS. Slices were conjugated with avidin-biotinylated horseradish peroxidase following the manufacturer’s instructions (ABC-Elite, Vector stains). Slices were then washed, and subsequently, biocytin-stained neurons were visualized under a reaction with 0.5 mg/ml DAB and 0.01% H2O2. When the neuronal processes were visible, the reaction was stopped by washing with PBS.

### Drugs and Chemicals

4-[3-[2-(Trifluoromethyl)-9H-thioxanthen-9-ylidene]propyl]-1-piperazineethanol dihydrochloride (flupenthixol dihydrochloride), [S-(R*,R*)]-[3-[[1-(3,4-Dichlorophenyl)ethyl]amino]-2-hydroxypropyl](cyclohexylmethyl) phosphinic acid (CGP 54626 hydrochloride), 8,8’-[Carbonylbis[imino-3,1-phenylenecarbonylimino(4-methyl-3,1-phenylene)carbonylimino]]bis-1,3,5-naphthalenetrisulfonic acid hexasodium salt (suramin hexasodium salt), N-(Piperidin-1-yl)-5-(4-iodophenyl)-1-(2,4-dichlorophenyl)-4-methyl-1H-pyrazole-3-carboxamide (AM251), D-(-)-2-Amino-5-phosphonopentanoic acid (D-AP5), 6-Cyano-7-nitroquinoxaline-2,3-dione disodium (CNQX disodium salt), (S)-(+)-a-Amino-4-carboxy-2-methylbenzeneacetic acid (LY367385), and 2-Methyl-6-(phenylethynyl)pyridine hydrochloride (MPEP hydrochloride), Octahydro-12-(hydroxymethyl)-2-imino-5,9:7,10a-dimethano-10aH-[1,3]dioxocino[6,5-d] pyrimidine-4,7,10,11,12-pentol (Tetrodotoxin: TTX) were purchased from Tocris Bioscience. Endo-(±)-α-(Hydroxymethyl)benzeneacetic acid 8-methyl-8-azabicyclo[3.2.1]oct-3-yl ester (atropine) and 8-Cyclopentyl-1,3-dimethylxanthine (CPT) were from Sigma. Picrotoxin from Indofine Chemical Company (Hillsborough, NJ). All other drugs were purchased from Sigma.

### Statistical analysis

Number of neurons was used as a sample size for electrophysiology comparisons and number of slices for Ca^2+^ signal comparisons. At least 2 mice per experimental group were used. Data are expressed as mean ± standard error of the mean (SEM). Data normality was tested using a Shapiro-Wilk test. Results were compared using a two-tailed Student’s t test (Paired, before-after stimulus-treatment; Unpaired between groups). A full report of the statistics used in every case is detailed in **Table S1**. Statistical differences were established with p < 0.05 (*), p < 0.01 (**) and p < 0.001 (***).

## ACKNOWLEDGMENTS

We would like to thank Dana Deters for technical support. We thank Carmen Nanclares, José Noriega, Francisco Emmanuel Labrada-Moncada, Julianna Goenaga, Carlos García, Pavan Guttipatti, Grace Gall and Jessica Neamtu helpful suggestions. We thank Justin Lines for providing Calsee and for helpful suggestions. We thank Guillermo Marques and Jason Mitchell at the University of Minnesota – University Imaging Centers for assistance using the Leica SP5 multiphoton upright microscope. We thank J. Chen (UCSD, USA) for providing IP_3_R_2_ mice. We thank the University of Minnesota Viral Vector and Cloning Core for production of some of the viral vectors used in this study. This work was supported by a postdoctoral fellowship from Basque Government, Spain, to AMB; grants from National Institutes of Health (NIH-MH R01MH119355; NIH-NINDS R01NS097312; and NIH-NIDA R01DA048822) to AA and grants from FEDER and ISCIII (AES2018-PI18/00513) and the Basque Government (PIBA19-0059) to SM, and ARSEP Foundation to SM and GM.

## AUTHOR CONTRIBUTIONS

A.M.B. performed experiments and analyzed data. L.B. and P.K. performed immunohistochemistry. P.K., A.A., G.M., C.M., S.M. and A.M.B. conceived the study and wrote the manuscript.

