## Supplemental Figures for "Spatial Organization of Neuron-Astrocyte Interactions in the Somatosensory Cortex"

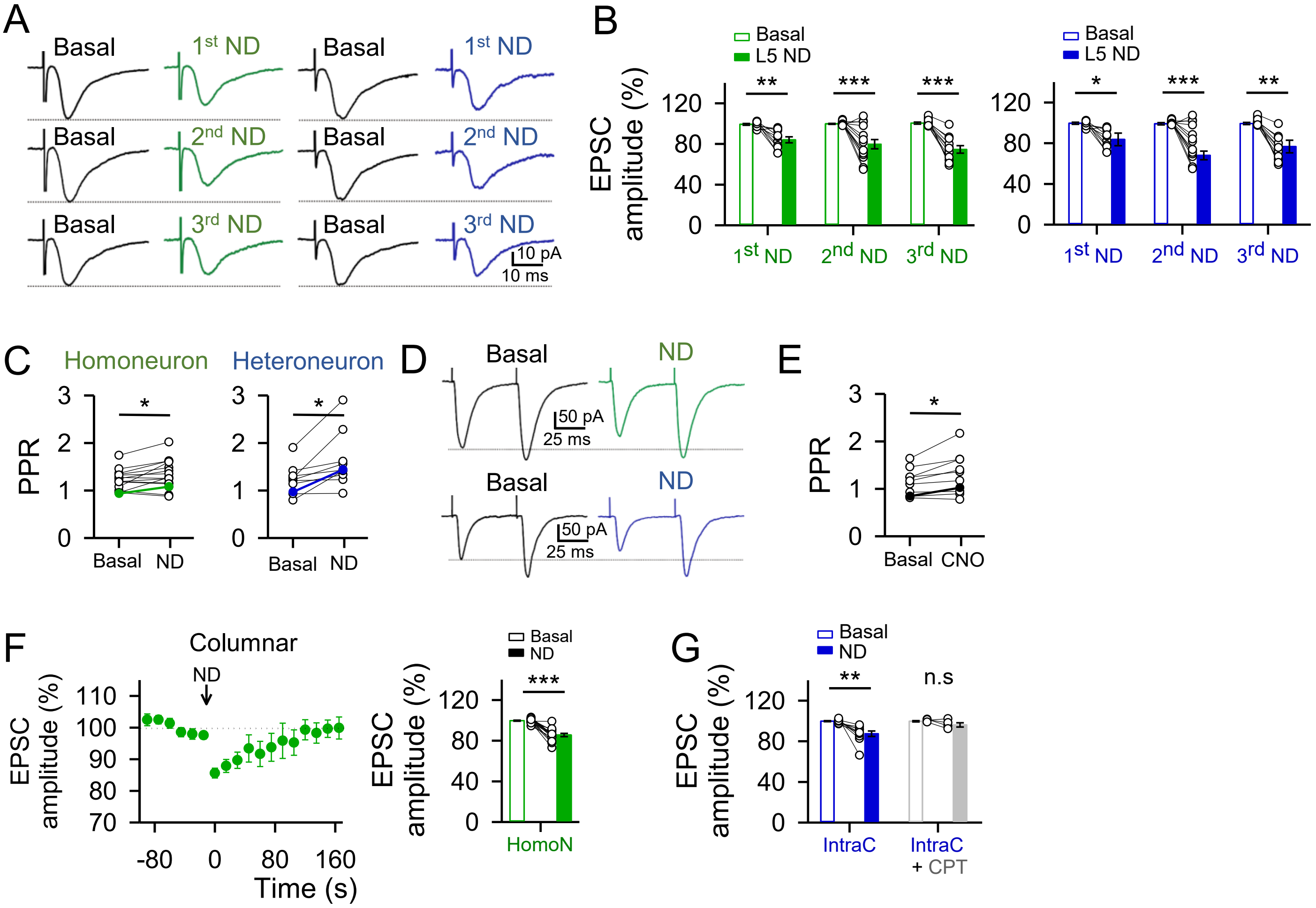


**Supplementary Figure 1. The effect of the endogenously mobilized eCBs is reliable and increases the pared pulse ratio.**

(**A**) Representative averaged EPSCs traces (n = 20 stimuli) before (control) and after three subsequent neuronal depolarizations (ND) in the homoneuron (left, green) and heteroneuron (right, blue) in wildtype mice. (**B**) Homoneuronal (left) and heteroneuronal (right) neuron relative EPSC amplitude before (basal) and after subsequent neuronal stimulations. Two-tailed Student's paired t test. (**C**) PPR = ($2^{nd}$ EPSC/$1^{st}$ EPSC) before (basal) and after ND in the homoneuron (left) and heteroneuron (right) from L5. Two-tailed Student's paired t test. (**D**) Representative pair of EPSCs traces separated by 50 ms and averaged 20 stimuli before (basal) and after ND in the homoneuron (top, green) and heteroneuron (bottom, blue). (**E**) PPR = ($2^{nd}$ EPSC/$1^{st}$ EPSC) before (basal) and after CNO application. Two-tailed Student's paired t test. (**F**) Left: EPSCs amplitude *versus* time before (basal) and after ND in the column homoneuron. Right: Relative changes in EPSC amplitude in the column homoneuron. (**G**) Relative changes in EPSC amplitude in control and with CPT (5 µM) in the intracolumn heteroneuron. Two-tailed Student's paired t test. Data are expressed as mean ± SEM, *$p < 0.05$, **$p < 0.01$, ***$p < 0.001$.

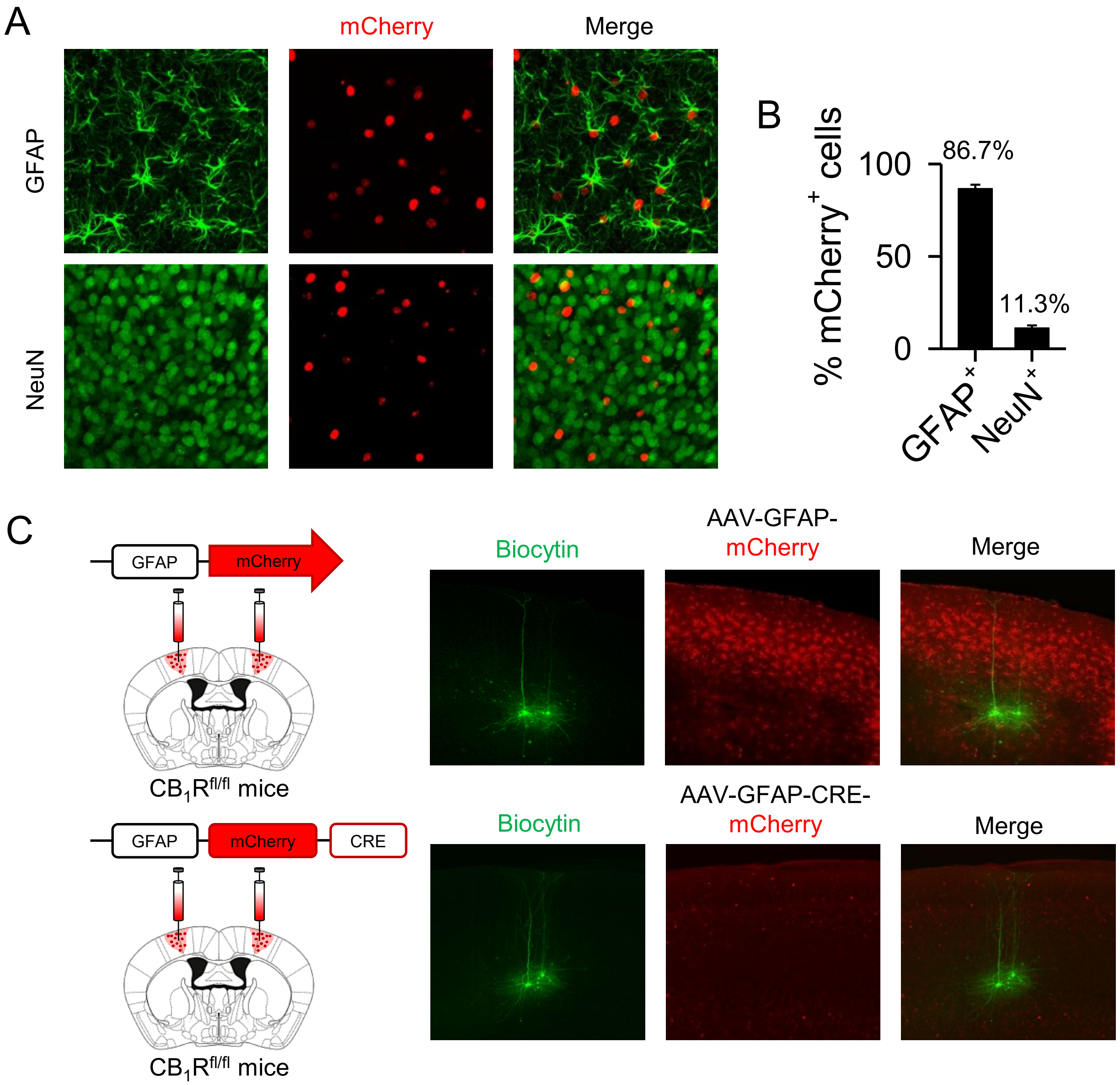


**Supplementary Figure 2. AAV8-GFAP-mCherry-Cre targets astrocytes from S1 cortex.**

(**A**) Immunohistochemistry images of mCherry$^{+}$ cells co-stained with GFAP and NeuN (n = 784 cells). From left to right: cell marker, mCherry, and merge. (**B**) Percent distribution of mCherry$^{+}$ cells. (**C**) Experimental scheme (left) and biocytin loading S1 L5 pyramidal neurons image in AAV8-GFAP-mCherry-Cre and AAV8-GFAP-mCherry injected mice.

**Statistical table**

| Figure | Comparison | Condition | Test | n | Mice | p values |
|---|---|---|---|---|---|---|
| Fig. 1E | Basal vs. after L5 ND | Homoneuron EPSCs amplitude (%) | Two-tailed Student's paired t-test | 36 | 18 | 3.141E-12 |
| Fig. 1E | Basal vs. after L5 ND | Homoneuron EPSCs amplitude (%) with AM251 | Two-tailed Student's paired t-test | 12 | 8 | 0.305 |
| Fig. 1E | Basal vs. after L5 ND | Heteroneuron EPSCs amplitude (%) | Two-tailed Student's paired t-test | 20 | 14 | 0.0000000567 |
| Fig. 1E | Basal vs. after L5 ND | Heteroneuron EPSCs amplitude (%) with AM251 | Two-tailed Student's paired t-test | 6 | 5 | 0.789 |
| Fig. 1G | Basal vs. after L4 ND | L4 homoneuron EPSCs amplitude (%) | Two-tailed Student's paired t-test | 8 | 3 | 0.008 |
| Fig. 1G | Basal vs. after L4 ND | L4 heteroneuron EPSCs amplitude (%) | Two-tailed Student's paired t-test | 8 | 3 | 0.0128 |
| Fig. 1I | Basal vs. after L2/3 ND | L2/3 homoneuron EPSCs amplitude (%) | Two-tailed Student's paired t-test | 14 | 7 | 0.00000441 |
| Fig. 1I | Basal vs. after L2/3 ND | L2/3 heteroneuron EPSCs amplitude (%) | Two-tailed Student's paired t-test | 14 | 6 | 0.0000543 |
| Fig. 2C | Basal vs. after L5 ND | Homoneuron EPSCs amplitude (%) in $aCB_1R$ mice | Two-tailed Student's paired t-test | 11 | 4 | 0.00226 |
| Fig. 2C | Basal vs. after L5 ND | Homoneuron EPSCs amplitude (%) in $aCB_1R^{-/-}$ mice | Two-tailed Student's paired t-test | 11 | 3 | 0.0000145 |
| Fig. 2C | Basal vs. after L5 ND | Heteroneuron EPSCs amplitude (%) in $aCB_1R$ mice | Two-tailed Student's paired t-test | 12 | 4 | 0.00000860 |
| Fig. 2C | Basal vs. after L5 ND | Heteroneuron EPSCs amplitude (%) in $aCB_1R^{-/-}$ mice | Two-tailed Student's paired t-test | 27 | 3 | 0.871 |
| Fig. 2E | Basal vs. after WIN application | $Ca^{2+}$ event probability in $aCB_1R$ mice | Two-tailed Student's paired t-test | 13 | 2 | 0.0000468 |
| Fig. 2E | Basal vs. after WIN application | $Ca^{2+}$ event probability in $aCB_1R^{-/-}$ mice | Two-tailed Student's paired t-test | 14 | 2 | 0.00765 |
| Fig. 2E | After WIN application | $Ca^{2+}$ event probability in $aCB_1R$ vs. $aCB_1R^{-/-}$ mice | Two-tailed Student's unpaired t-test | 13 vs. 14 | 2 vs. 2 | 0.005 |
| Fig. 3B | Basal vs. after L5 ND | $Ca^{2+}$ event probability | Two-tailed Student's paired t-test | 5 | 4 | 0.000755 |
| Fig. 3B | Basal vs. after L5 ND | $Ca^{2+}$ event probability with AM251 | Two-tailed Student's paired t-test | 5 | 4 | 0.422 |
| Fig. 3B | Basal vs. after L5 ND | $Ca^{2+}$ event probability in $IP_3R_2^{-/-}$ mice | Two-tailed Student's paired t-test | 11 | 3 | 0.889 |
| Fig. 3B | Basal vs. after L5 ND | $Ca^{2+}$ event probability in $IP_3R_2^{-/-}$ mice with AM251 | Two-tailed Student's paired t-test | 6 | 2 | 0.749 |
| Fig. 3C | Basal vs. after L5 ND | Homoneuron EPSCs amplitude (%) | Two-tailed Student's paired t-test | 36 | 18 | 3.141E-12 |
| Fig. 3C | Basal vs. after L5 ND | Homoneuron EPSCs amplitude (%) in $IP_3R_2^{-/-}$ mice | Two-tailed Student's paired t-test | 13 | 3 | 0.00933 |
| Fig. 3C | Basal vs. after L5 ND | Homoneuron EPSCs amplitude (%) in WT with CPT | Two-tailed Student's paired t-test | 8 | 2 | 0.00416 |
| Fig. 3C | Basal vs. after L5 ND | Heteroneuron EPSCs amplitude (%) | Two-tailed Student's paired t-test | 20 | 14 | 0.0000000567 |
| Fig. 3C | Basal vs. after L5 ND | Heteroneuron EPSCs amplitude (%) in $IP_3R_2^{-/-}$ mice | Two-tailed Student's paired t-test | 16 | 3 | 0.0796 |
| Fig. 3C | Basal vs. after L5 ND | Heteroneuron EPSCs amplitude (%) in WT with CPT | Two-tailed Student's paired t-test | 7 | 3 | 0.171 |
| Fig. 3D | Basal vs. after L2/3 ND | L2/3 homoneuron EPSCs amplitude (%) | Two-tailed Student's paired t-test | 14 | 7 | 0.00000441 |
| Fig. 3D | Basal vs. after L2/3 ND | L2/3 homoneuron EPSCs amplitude (%) with CPT | Two-tailed Student's paired t-test | 5 | 3 | 0.0294 |
| Fig. 3D | Basal vs. after L2/3 ND | L2/3 heteroneuron EPSCs amplitude (%) | Two-tailed Student's paired t-test | 14 | 6 | 0.0000543 |
| Fig. 3D | Basal vs. after L2/3 ND | L2/3 heteroneuron EPSCs amplitude (%) with CPT | Two-tailed Student's paired t-test | 7 | 3 | 0.139 |
| Fig. 3D | Basal vs. after L4 ND | L4 homoneuron EPSCs amplitude (%) | Two-tailed Student's paired t-test | 8 | 3 | 0.008 |
| Fig. 3D | Basal vs. after L4 ND | L4 homoneuron EPSCs amplitude (%) with CPT | Two-tailed Student's paired t-test | 4 | 2 | 0.042 |
| Fig. 3D | Basal vs. after L4 ND | L4 heteroneuron EPSCs amplitude (%) | Two-tailed Student's paired t-test | 8 | 3 | 0.0128 |
| Fig. 3D | Basal vs. after L4 ND | L4 heteroneuron EPSCs amplitude (%) with CPT | Two-tailed Student's paired t-test | 4 | 2 | 0.680 |
| Fig. 3F | Basal vs. after CNO application | EPSCs amplitude (%) in mice injected with DREADDs | Two-tailed Student's paired t-test | 11 | 2 | 0.00138 |
| Fig. 3F | Basal vs. after CNO application | EPSCs amplitude (%) with CPT in mice injected with DREADDs | Two-tailed Student's paired t-test | 7 | 2 | 0.716 |
| Fig. 3F | Basal vs. after CNO application | EPSCs amplitude (%) in mice injected with mCherry | Two-tailed Student's paired t-test | 11 | 2 | 0.309 |
| Fig. 3G | Basal vs. after CNO application | $Ca^{2+}$ event probability in mice injected with DREADDs | Two-tailed Student's paired t-test | 14 | 3 | 0.0000392 |
| Fig. 3G | Basal vs. after CNO application | $Ca^{2+}$ event probability with CPT in mice injected with DREADDs | Two-tailed Student's paired t-test | 7 | 3 | 0.00956 |
| Fig. 3G | Basal vs. after CNO application | $Ca^{2+}$ event probability in mice injected with mCherry | Two-tailed Student's paired t-test | 8 | 2 | 0.158 |
| Fig. 4B | Basal vs. after L4 ND | Heteroneuron intracolumnar EPSCs amplitude (%) | Two-tailed Student's paired t-test | 10 | 6 | 0.0000918 |
| Fig. 4C | Basal vs. after L4 ND | Heteroneuron intercolumnar EPSCs amplitude (%) | Two-tailed Student's paired t-test | 11 | 3 | 0.0797 |
| Fig. 5B | Basal vs. after L4 ND | L2/3 heteroneuron EPSCs amplitude (%) | Two-tailed Student's paired t-test | 5 | 4 | 0.00354 |
| Fig. 5B | Basal vs. after L2/3 ND | L4 heteroneuron EPSCs amplitude (%) | Two-tailed Student's paired t-test | 9 | 4 | 0.0625 |
| Fig. 5D | Basal vs. after L5 ND | L2/3 heteroneuron EPSCs amplitude (%) | Two-tailed Student's paired t-test | 4 | 2 | 0.0195 |
| Fig. 5D | Basal vs. after L2/3 ND | L5 heteroneuron EPSCs amplitude (%) | Two-tailed Student's paired t-test | 8 | 2 | 0.147 |
| Fig. 5F | Basal vs. after L4 ND | L5 heteroneuron EPSCs amplitude (%) | Two-tailed Student's paired t-test | 4 | 3 | 0.0320 |
| Fig. 5F | Basal vs. after L5 ND | L4 heteroneuron EPSCs amplitude (%) | Two-tailed Student's paired t-test | 14 | 4 | 0.0552 |
| Fig. 6B | Basal vs. after L2/3 ND | L2/3 astrocytic $Ca^{2+}$ event probability | Two-tailed Student's paired t-test | 5 | 4 | 0.0303 |
| Fig. 6B | Basal vs. after L2/3 ND | L2/3 astrocytic $Ca^{2+}$ event probability with AM251 | Two-tailed Student's paired t-test | 5 | 4 | 0.136 |
| Fig. 6C | Basal vs. after L4 ND | L4 astrocytic $Ca^{2+}$ event probability | Two-tailed Student's paired t-test | 4 | 3 | 0.00282 |
| Fig. 6C | Basal vs. after L4 ND | L4 astrocytic $Ca^{2+}$ event probability with AM251 | Two-tailed Student's paired t-test | 4 | 3 | 0.124 |
| Fig. 6D | Basal vs. after L5 ND | L5 astrocytic $Ca^{2+}$ event probability | Two-tailed Student's paired t-test | 5 | 4 | 0.000755 |
| Fig. 6D | Basal vs. after L5 ND | L5 astrocytic $Ca^{2+}$ event probability with AM251 | Two-tailed Student's paired t-test | 5 | 4 | 0.422 |
| Fig. 6E | Basal vs. after L2/3 ND | L2/3 astrocytic $Ca^{2+}$ event probability | Two-tailed Student's paired t-test | 10 | 7 | 0.000852 |
| Fig. 6E | Basal vs. after L2/3 ND | L4 astrocytic $Ca^{2+}$ event probability | Two-tailed Student's paired t-test | 5 | 3 | 0.148 |
| Fig. 6E | Basal vs. after L2/3 ND | L5 astrocytic $Ca^{2+}$ event probability | Two-tailed Student's paired t-test | 6 | 4 | 0.309 |
| Fig. 6F | Basal vs. after L4 ND | L2/3 astrocytic $Ca^{2+}$ event probability | Two-tailed Student's paired t-test | 6 | 3 | 0.031 |
| Fig. 6F | Basal vs. after L4 ND | L4 astrocytic $Ca^{2+}$ event probability | Two-tailed Student's paired t-test | 8 | 5 | 0.0109 |
| Fig. 6F | Basal vs. after L4 ND | L5 astrocytic $Ca^{2+}$ event probability | Two-tailed Student's paired t-test | 7 | 3 | 0.0153 |
| Fig. 6G | Basal vs. after L5 ND | L2/3 astrocytic $Ca^{2+}$ event probability | Two-tailed Student's paired t-test | 8 | 4 | 0.008 |
| Fig. 6G | Basal vs. after L5 ND | L4 astrocytic $Ca^{2+}$ event probability | Two-tailed Student's paired t-test | 8 | 4 | 0.101 |
| Fig. 6G | Basal vs. after L5 ND | L5 astrocytic $Ca^{2+}$ event probability | Two-tailed Student's paired t-test | 13 | 7 | 0.001 |
| Fig. S1B | Basal vs. after 1º ND | Homoneuron EPSCs amplitude (%) | Two-tailed Student's paired t-test | 10 | 8 | 0.00181 |
| Fig. S1B | Basal vs. after 2º ND | Homoneuron EPSCs amplitude (%) | Two-tailed Student's paired t-test | 14 | 8 | 0.000864 |
| Fig. S1B | Basal vs. after 3º ND | Homoneuron EPSCs amplitude (%) | Two-tailed Student's paired t-test | 11 | 8 | 0.0000232 |
| Fig. S1B | Basal vs. after 1º ND | Heteroneuron EPSCs amplitude (%) | Two-tailed Student's paired t-test | 9 | 8 | 0.0294 |
| Fig. S1B | Basal vs. after 2º ND | Heteroneuron EPSCs amplitude (%) | Two-tailed Student's paired t-test | 11 | 8 | 0.0000283 |
| Fig. S1B | Basal vs. after 3º ND | Heteroneuron EPSCs amplitude (%) | Two-tailed Student's paired t-test | 9 | 8 | 0.00607 |
| Fig. S1C | Basal vs. after ND | Homoneuron PPR | Two-tailed Student's paired t-test | 15 | 6 | 0.0165 |
| Fig. S1C | Basal vs. after ND | Heteroneuron PPR | Two-tailed Student's paired t-test | 9 | 6 | 0.0180 |
| Fig. S1E | Basal vs. after CNO application | PPR | Two-tailed Student's paired t-test | 11 | 2 | 0.0166 |
| Fig. S1F | Basal vs. after L4 ND | Homoneuron columnar EPSCs amplitude (%) | Two-tailed Student's paired t-test | 16 | 7 | 0.000000292 |
| Fig. S1G | Basal vs. after L4 ND | Heteroneuron intracolumnar EPSCs amplitude (%) | Two-tailed Student's paired t-test | 10 | 6 | 0.0000918 |
| Fig. S1G | Basal vs. after L4 ND | Heteroneuron intracolumnar EPSCs amplitude (%) with CPT | Two-tailed Student's paired t-test | 4 | 3 | 0.153 |
| Fig. S2B | | % $mCherry^+$ cells $GFAP^+$ | | 9 | 2 | |
| Fig. S2B | | % $mCherry^+$ cells $NeuN^+$ | | 11 | 2 | |

**Supplementary Table 1**. **Full report of Two-tailed Student's paired t-test and Unpaired t-test values.**

Full report of comparison, condition, test, n and number of mice used and their p values.
